# Exploiting flow cytometry for the unbiased quantification of protein inclusions in *Caenorhabditis elegans*

**DOI:** 10.1101/2021.08.29.458141

**Authors:** Kristian Claesson, Yee Lian Chew, Heath Ecroyd

## Abstract

The aggregation of proteins into inclusions or plaques is a prominent hallmark of a diverse range of pathologies including neurodegenerative diseases. The quantification of such inclusions in *Caenorhabditis elegans* models of aggregation is usually achieved by fluorescence microscopy or other techniques involving biochemical fractionation of worm lysates. Here, we describe a simple and rapid flow cytometry-based approach that allows fluorescently-tagged inclusions to be enumerated in whole worm lysate in a quantitative and unbiased fashion. We demonstrate that this technique is applicable to multiple *C. elegans* models of aggregation and importantly, can be used to monitor the dynamics of inclusion formation in response to heat shock and during aging. This includes the characterisation of physicochemical properties of inclusions, such as their size, which may reveal how aggregate formation is distinct in different tissues or at different stages of pathology or aging. This new method can be used as a powerful technique for the medium- to high-throughput quantification of inclusions in future studies of genetic or chemical modulators of aggregation in *C. elegans*.

## Introduction

A common pathological hallmark of many diseases, including the neurodegenerative disorders Alzheimer’s disease, Huntington’s disease and amyotrophic lateral sclerosis (ALS), is the aberrant aggregation and deposition of proteins, in the form of cytoplasmic and nuclear inclusions within cells, or plaques in the extracellular space (Aguzzi & O’Connor 2010). For the vast majority of pathologies associated with protein aggregation (proteinopathies), there are currently no effective treatments for preventing the onset or progression of disease. Thus, there is a pressing need for the development of new pharmacological approaches that target key components of this process. Since the formation of insoluble proteinaceous inclusion bodies is a common feature among proteinopathies, it is thought that preventing this process is likely to be a beneficial strategy for halting the progression of these diseases (Bose & Cho 2017). Thus, multiple studies have focused on high-throughput screening for genetic or chemical modulators of protein aggregation in various model systems (Cockburn et al. 2011; Ikenaka et al. 2019; Sarkany et al. 2019; Silva et al. 2011). These high-throughput screens often rely on having a quantifiable readout of aggregation dynamics that can be determined rapidly and at low-cost.

The translucent, soil-dwelling nematode *Caenorhabditis elegans* (*C. elegans*) is a powerful model organism for the study of molecular processes (including those perturbed in disease), as many genes and complex biochemical pathways are conserved with mammals (Alexander et al. 2014). In addition, these worms have a short reproductive cycle and are easy to manipulate genetically. Consequently, they have been investigated extensively in forward and reverse genetic studies to identify genes that modulate biochemical processes including those involved in protein aggregation and neurodegeneration (Kraemer et al. 2006; Nollen et al. 2004; van Ham et al. 2008). They are also highly amenable to screens of pharmacological compounds and are often used as an initial model system for potential therapeutics before conducting rodent-based studies (Chen et al. 2015; Ikenaka et al. 2019). A common approach for studying neurodegenerative diseases in *C. elegans* is to utilise strains that express specific disease-associated, aggregation-prone proteins fused with a fluorescent tag for visualisation within worms. For example, there are several established *C. elegans* strains that attempt to model ALS by expressing the ALS-associated protein superoxide dismutase 1 (SOD1). Worms that pan-neuronally express SOD1 with the destabilising disease-associated mutation, G85R, form inclusions in neuronal cell bodies, and have motility defects and a shortened lifespan (Wang et al. 2009).

A fundamental component of studying the aggregation of proteins in cells and organisms, and identifying modulators of this process, is the capacity to quantify the number and type of inclusions that are formed. The quantification of aggregation in *C. elegans* often requires laborious manual counting of inclusions from images generated through fluorescence microscopy, or other biochemical methods, such as immunoblotting or filter trap assays, which rely on quantifying the amount of insoluble protein in the lysate (Sin et al. 2018; Walther et al. 2017). Recently, there have been several advances in automated imaging methods in an attempt to increase the throughput of microscopy-based experiments (Cornaglia et al. 2016; Gosai et al. 2010; Mondal et al. 2016). However, there are inherent limitations with these approaches, such as difficulty in distinguishing closely positioned inclusions and smaller aggregates (e.g. inclusions that form within neuronal cell bodies in worms).

The recent application of flow cytometry-based approaches to quantifying aggregation in cell-based studies has shown promise in that it circumvents some of the limitations associated with using fluorescence microscopy to identify and enumerate inclusions. One such technique is the pulse shape analysis (PulSA) of whole cells (Ramdzan et al. 2012). This technique can be used to identify cultured cells that contain protein aggregates on the basis that the presence of large fluorescent inclusions within cells alters their pulse profile. More recently, flow cytometry has been used to directly quantify the number of inclusions in cell lysates. Shiber et al. (2014) described a technique using flow cytometry to quantify aggregation of fluorescent proteins in yeast cells by lysing cells and normalising the detected inclusions to the protein content of the lysate. Subsequently, a similar approach known as FloIT (flow cytometric analysis of inclusions and trafficking) was developed and applied to mammalian cells (Whiten et al. 2016). This method involves lysing cells with a mild detergent and normalising detected fluorescent inclusions to the number of nuclei (and thus, the relative number of cells) in the sample. This technique also has the advantage of allowing for the quantification of nuclear flux of fluorescently-tagged proteins. The application of flow cytometric analyses to quantify inclusions has been demonstrated to be a powerful tool for cell-based studies as they are unbiased and can be applied to high-throughput screens (McMahon et al. 2021; Whiten et al. 2016). However, a method exploiting flow cytometry to quantify inclusions formed in *C. elegans* has not yet been described. Thus, in this study we sought to devise an efficient, unbiased, and quantitative method to enumerate protein inclusions in whole worm lysate using a flow cytometry-based approach.

## Results

As a proof of concept for applying a flow cytometric approach for the quantification of aggregation in nematodes, we first utilised a recently developed *C. elegans* model of Huntington’s disease (Lee et al. 2017). In this model, the first 513 residues of the N-terminal fragment of the Huntingtin protein (Htt) are followed by a polyglutamine (polyQ) tract containing either 15 (HttQ15) or 128 (HttQ128) glutamine residues, fused to a yellow fluorescent protein (YFP) tag at the C-terminus. The expression of these Htt constructs is controlled by the myosin heavy chain promoter, *unc-54*, which drives expression in the body-wall muscle cells of *C. elegans*. In Huntington’s disease, polyQ expansions within Htt increase its aggregation propensity in a polyQ length-dependent fashion (Imarisio et al. 2008). As previously reported (Lee et al. 2017), the HttQ15 expressing worm strain exhibited diffuse fluorescence in the muscle tissue, whereas bright fluorescent foci were present in the HttQ128 strain, indicating the aggregation of HttQ128 into inclusion bodies **(Fig. 1A)**.

**Figure 1:**
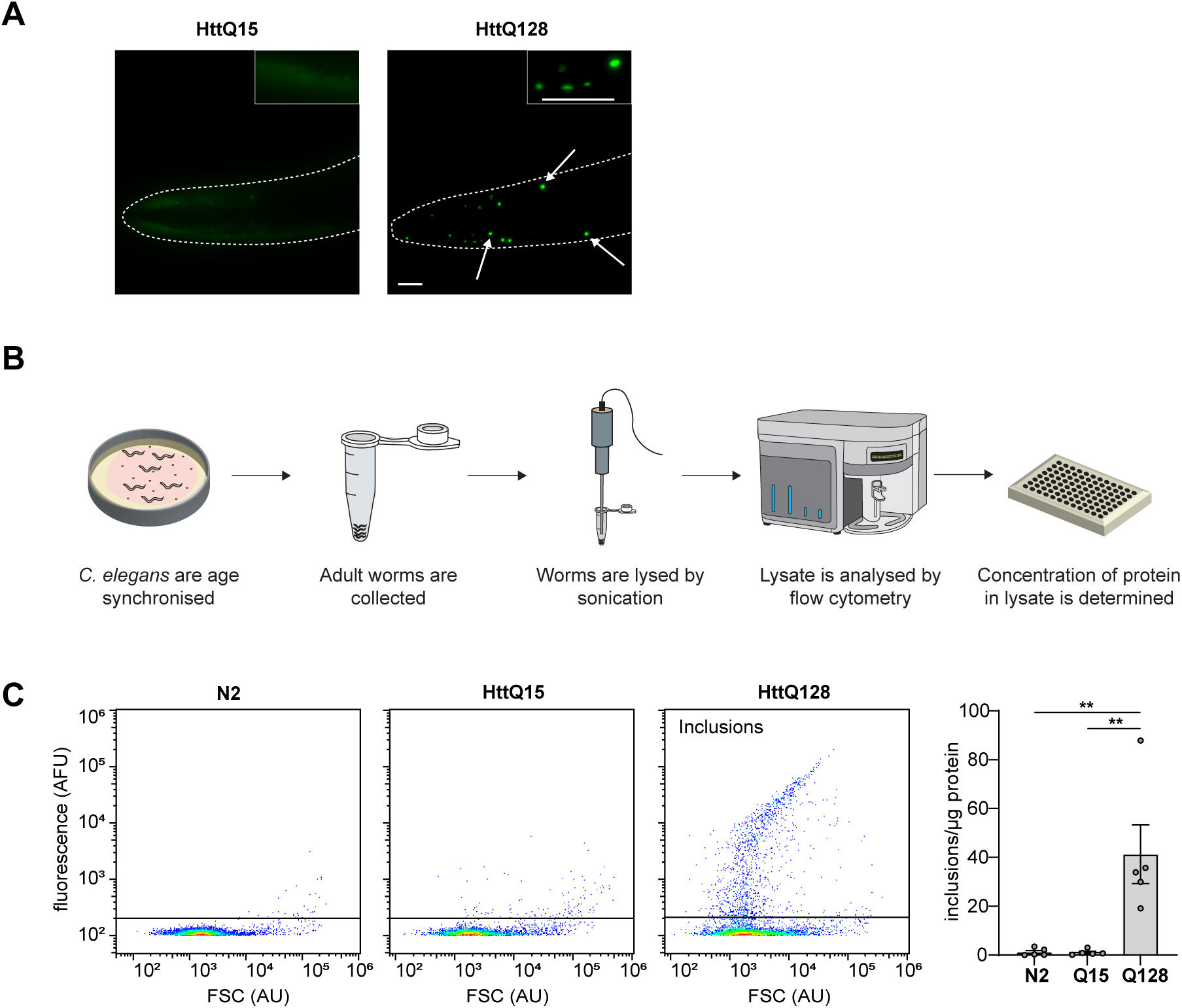
Flow cytometric analysis of worm lysate is a viable approach for quantifying fluorescently-tagged protein inclusions in *C. elegans*. **(a)** Epifluorescence microscopy images of day 1 adult worms expressing HttQ15 (*left*) or HttQ128 (*righ*t) in body-wall muscle cells. Insets show digitally magnified regions of the respective images. Arrows indicate examples of protein inclusions. Scale bars are 20 μm. **(b)** A schematic of the flow cytometry-based approach for the detection and quantification of fluorescent protein inclusions in *C. elegans* lysate. **(c)** Two parameter, pseudo-colour flow cytometry dot plots of forward scatter (FSC) versus YFP-derived fluorescence of lysates from N2, Htt15Q and Htt128Q worm strains allowing identification of inclusions (indicated). Axes are in logarithmic scale. Histogram (right) shows quantification of number of inclusions per μg of protein in these strains. Data are presented as the mean ± S.E.M with data points from 5 independent experiments shown. Statistical significance between group means in the data was determined using a one-way ANOVA followed by a Tukey’s post-hoc test. ** denotes a statistically significant difference between indicated samples (p < 0.01).

As an approach to quantify the aggregation profile in these strains, lysate from ~500 age-synchronised day 1 adult worms was generated by sonication and immediately analysed by flow cytometry **(Fig. 1B)**. Lysates from worms expressing YFP-tagged Htt and the non-fluorescent wildtype (N2) *C. elegans* strain were analysed by plotting forward scatter (FSC) against YFP fluorescence such that the relative fluorescence intensity of insoluble particles (events) in the lysate was measured. In the N2 control lysate, a population of particles was detected (spanning a range of FSC values) that represented cellular debris and organelles **(Fig. 1C)**. As expected, these particles did not exhibit any significant YFP fluorescence. The lysate from HttQ15 worms had a very similar profile to the N2 lysate, indicating that the majority of the fluorescent protein in the sample was soluble and not detectable by flow cytometry **(Fig. 1C)**. However, in the HttQ128 lysate, a clear population of YFP positive particles were detected, representing insoluble fluorescent inclusions **(Fig. 1C)**. This result established that flow cytometric analysis of *C. elegans* lysate could indeed detect protein inclusions. By normalising the number of YFP positive events detected in the ‘inclusions’ gate to the total concentration of protein in the lysate, the HttQ128 sample was found to contain 41.3 ± 12.0 inclusions/μg of protein in the lysate, which was significantly more (p < 0.01) than that detected in the lysate from HttQ15 worms (1.1 ± 0.5 inclusions/μg of protein in the lysate).

Next, we wanted to further explore the applicability of using this flow cytometric analysis of worm lysate technique on other model strains relevant to neurodegenerative disease. To achieve this, we utilised strains generated to model ALS pathology by expressing disease-associated isoforms of SOD1. The strains that we exploited included one that expresses YFP-tagged human SOD1 with the disease-linked G85R mutation in all neurons, (Wang et al. 2009) and another in which SOD1 carrying a 127X (G127insTGGGstop) frameshift mutation was expressed in body-wall muscle cells (Gidalevitz et al. 2009). Both of these mutations in SOD1 make the protein more aggregation-prone (Crown et al. 2020; Lang et al. 2015). Corresponding strains that express wildtype human SOD1 (WT SOD1) in neurons or muscle were used as controls in these experiments as WT SOD1 is significantly less aggregation-prone than the G85R and 127X mutants (Gidalevitz et al. 2009; Wang et al. 2009). In both strains expressing disease-associated SOD1 mutations, bright fluorescent foci were identified either in neuronal cell bodies (G85R) or body-wall muscle cells (127X) **(Fig. 2A)**. By fractionating whole worm lysate and analysing it by immunoblotting with anti-GFP antibodies, it was confirmed that the fluorescent foci observed in these strains represented NP-40-insoluble SOD1 protein inclusions, evidenced by the prominent band detected at ~ 50 kDa corresponding to the molecular mass of YFP-tagged SOD1 (Prudencio & Borchelt 2011), and degraded products of lower molecular weight **(Fig. 2B & S2)**. In contrast, both WT SOD1 expressing control strains had diffuse fluorescence in these tissues **(Fig. 2A)**, which correlated to less insoluble material being detected by immunoblotting the lysate from these strains **(Fig. 2B)**. Flow cytometry of the cell lysates from these worm strains allowed clear identification of insoluble inclusions in the disease-related mutant strains **(Fig. 2C)**. Using the flow cytometry data to quantify these inclusions, it was found that 174.8 ± 19.0 inclusions/μg of protein were detected in the G85R sample, which was significantly higher (p < 0.001) than the number of inclusions detected in the neural-expressing WT SOD1 sample (8.2 ± 5.3 inclusions/μg of protein). Similarly, significantly more inclusions (p < 0.001) were identified in the lysate from the 127X strain (1369 ± 81.5 inclusions/μg of protein) compared to the lysate from the control strain expressing WT SOD1 in muscle tissue (132.1 ± 23.2 inclusions/μg of protein).

**Figure 2:**
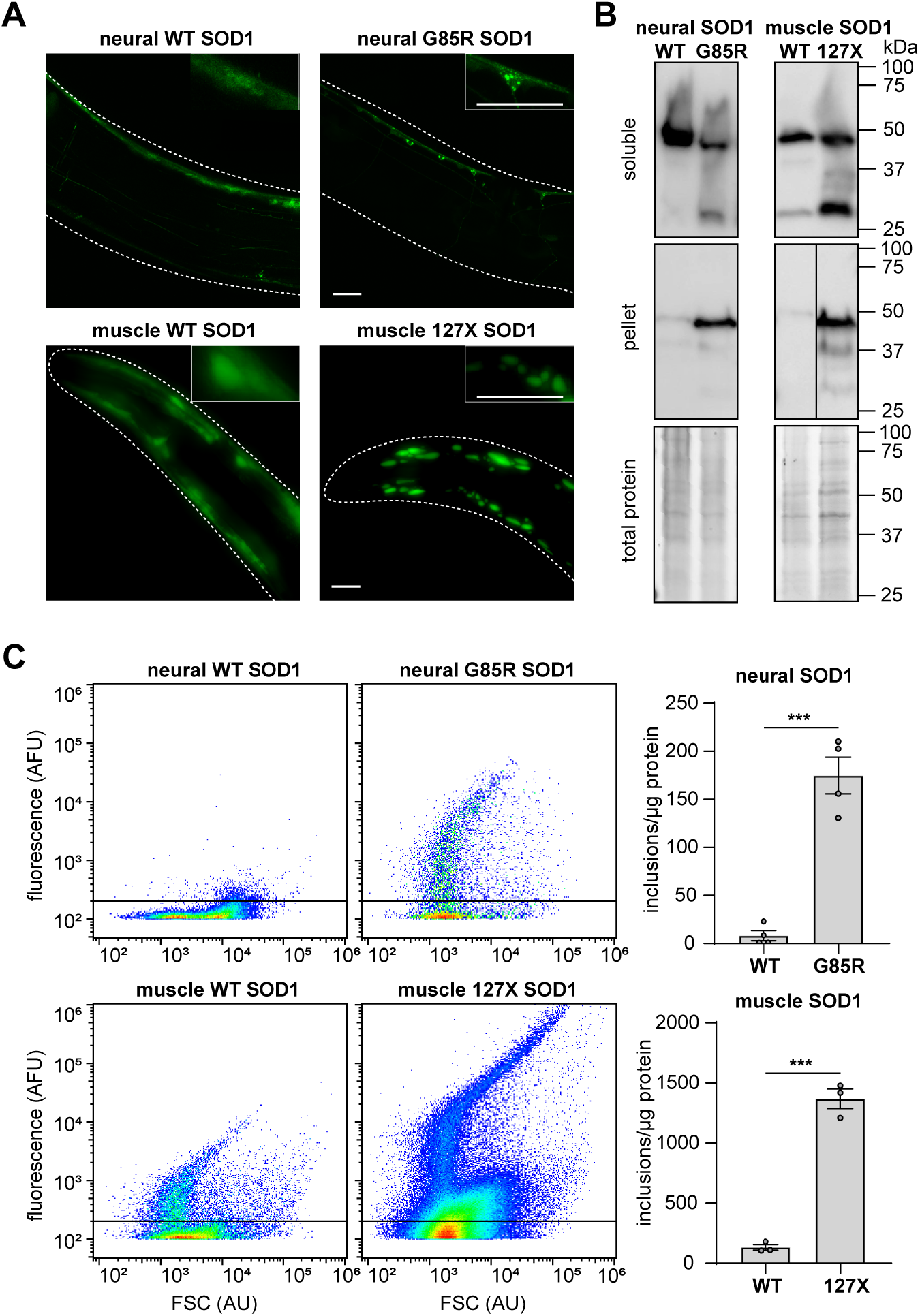
Flow cytometric analysis of worm lysate to quantify inclusions is a technique applicable to *C. elegans* models of ALS. **(a)** Strains expressing YFP-tagged human SOD1 mutants in all neurons (*top*) or in body-wall muscle cells (*bottom*) were imaged to visualise SOD1::YFP. Insets in the images show digitally magnified regions of the respective images. Scale bars are 20 μm. **(b)** Worm lysates were separated into soluble and insoluble fractions by centrifugation and analysed by immunoblotting. An anti-GFP antibody was used to detect the YFP-tagged SOD1 isoforms (~50 kDa). The line separating WT and 127X in the pellet fraction of the muscle SOD1 samples indicates these samples were run on the same SDS PAGE gel but were not in adjacent lanes. The relative amount of total protein in each lysate is shown. **(c)** Two parameter, pseudo-colour flow cytometry dot plots of forward scatter (FSC) versus YFP-derived fluorescence of lysates from each SOD1-expressing strain. Axes are in logarithmic scale. Quantification of the number of inclusions per μg of protein in SOD1-expressing strains is shown to the right of the dot plots. Data points in graphs represent values from either 4 (*top*) or 3 (*bottom*) independent replicates. Data are shown as the mean ± S.E.M. A student’s t-test was used to determine statistical significance between group means in the data. *** denotes a statistically significant difference between indicated samples (p < 0.001).

Having established that fluorescent inclusions could be detected in the lysate of multiple *C. elegans* models of neurodegenerative disease using flow cytometry, we next sought to determine whether this technique could detect dynamic changes in inclusion formation within a worm strain. To do so, and to further demonstrate the broad applicability of this approach, we assayed worms that express EGFP-tagged firefly luciferase (Fluc) variants in either neurons or body-wall muscle cells, as these have previously been shown to be sensitive to modulation of protein homeostasis (Gupta et al. 2011). The variants used in this study were the wildtype form of Fluc and a more aggregation-prone double mutant form (R188Q, R261Q), referred to here as FlucWT and FlucDM, respectively. Consistent with the previous report by Gupta et al. (2011), using fluorescence microscopy we observed that FlucWT did not form inclusions in neural or muscle cells in young adult worms, and the protein remained diffuse after acute heat shock (35°C for 1 h) **(Fig. 3A)**. In contrast, FlucDM formed inclusions in both neurons and muscle cells in young adult worms. Acute heat shock induced an immediate and robust increase in inclusion formation of FlucDM. A short recovery period (5 h at 20°C) was sufficient to significantly reduce the aggregate load in both FlucDM strains, further corroborating the findings of Gupta et al. (2011).

**Figure 3:**
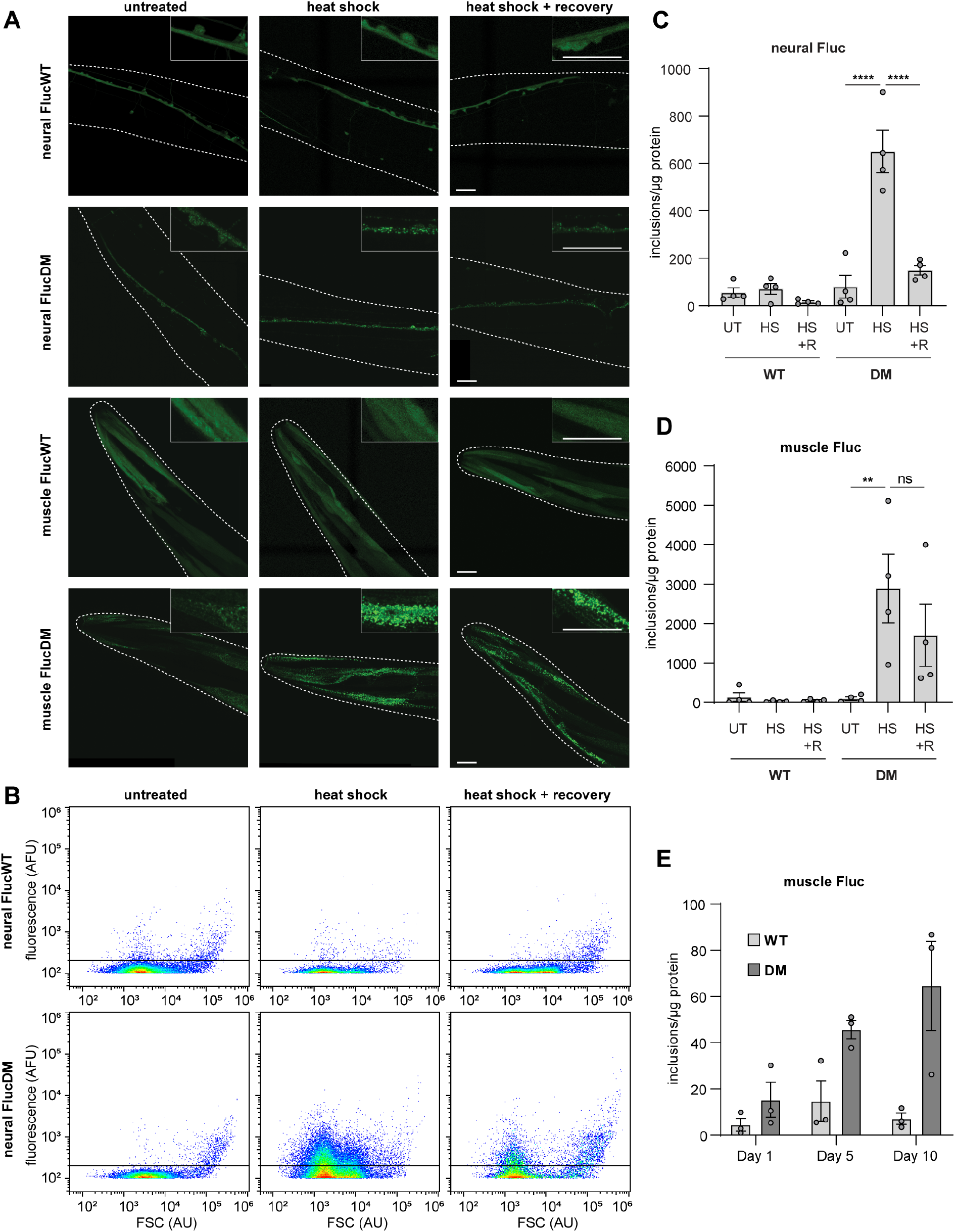
Flow cytometric analysis of worm lysates can be used to monitor changes in inclusion formation in response to stress and aging. **(a)** Confocal microscopy images of day 1 adult worms expressing wild type (WT) or double mutant (DM) Fluc-EGFP in neurons (showing ventral nerve cord) or body-wall muscle cells under normal conditions (untreated), after heat shock at 35°C for 1 h, or with the heat shock treatment followed by a 5 h recovery period at 20°C. Insets in the images show digitally magnified regions of the respective images. Scale bars are 20 μm. Note that worms are twisted. **(b)** Two parameter, pseudo-colour flow cytometry dot plots of forward scatter (FSC) versus EGFP-derived fluorescence of (*top*) neural FlucWT or (*bottom*) neural FlucDM worm lysates, following the same treatments as described in (a). Axes are in logarithmic scale. **(c)-(e)** Quantification of the number of inclusions per μg of protein in neural and muscle Fluc lysates by flow cytometry. Data points in graphs represent values from either 4 (*c and d*) or 3 (*e*) independent replicates. Data are represented as the mean ± S.E.M. Significant differences between group means in the data were determined using a two-way ANOVA followed by a Tukey’s post hoc test. ** and **** denote statistically significant differences between indicated samples (p < 0.01 and p < 0.0001, respectively).

By using our flow cytometric method we were able to detect and quantify changes in Fluc aggregation in these *C. elegans* strains **(Fig. 3B)**. There was no notable increase in EGFP positive particles detected in response to heat shock in either the neural or muscle FlucWT strains (**Fig. 3C & D**). In contrast, in both muscle and neural-expressing FlucDM strains there was a significant increase in the number of inclusions detected by flow cytometry in response to heat shock, compared with untreated controls. The number of inclusions decreased following recovery from heat stress, and this reached statistical significance for neural FlucDM worms **(Fig 3C & D)**. Next, we applied the flow cytometry technique to worms of different ages, including days 1, 5 and 10 of adulthood, using the strains expressing the Fluc variants in muscle cells. We found that our flow cytometric method was able to discern the age-associated increase in aggregation of FlucDM, but not FlucWT, in muscle as had been reported previously (Gupta et al. 2011; Walther et al. 2017) **(Fig. 3E)**.

Finally, we exploited the capacity to determine the size of detected particles using the FSC signal of the flow cytometer in order to further characterise the inclusions present in the worm lysates. By calibrating the FSC signal using size calibration microspheres **(Fig. S1)**, the size distribution of inclusions detected in different transgenic lines could be determined **(Fig. 4)**. FlucDM inclusions were similar in size, with the majority around 1 μm in diameter, in both neurons and body-wall muscle cells. This corresponded to their approximate size *in vivo* when observed by confocal microscopy (see Fig. 3A). HttQ128 and muscle 127X SOD1 inclusions encompassed a broader range of sizes and tended to be larger, again in accord with their larger appearance in the fluorescence micrographs of these worms (see Fig. 1A & 2A). Together, these data demonstrate the versatility of this flow cytometry approach in being able to enumerate and characterise physicochemical properties of inclusions in worms in a rapid and non-subjective manner.

**Figure 4:**
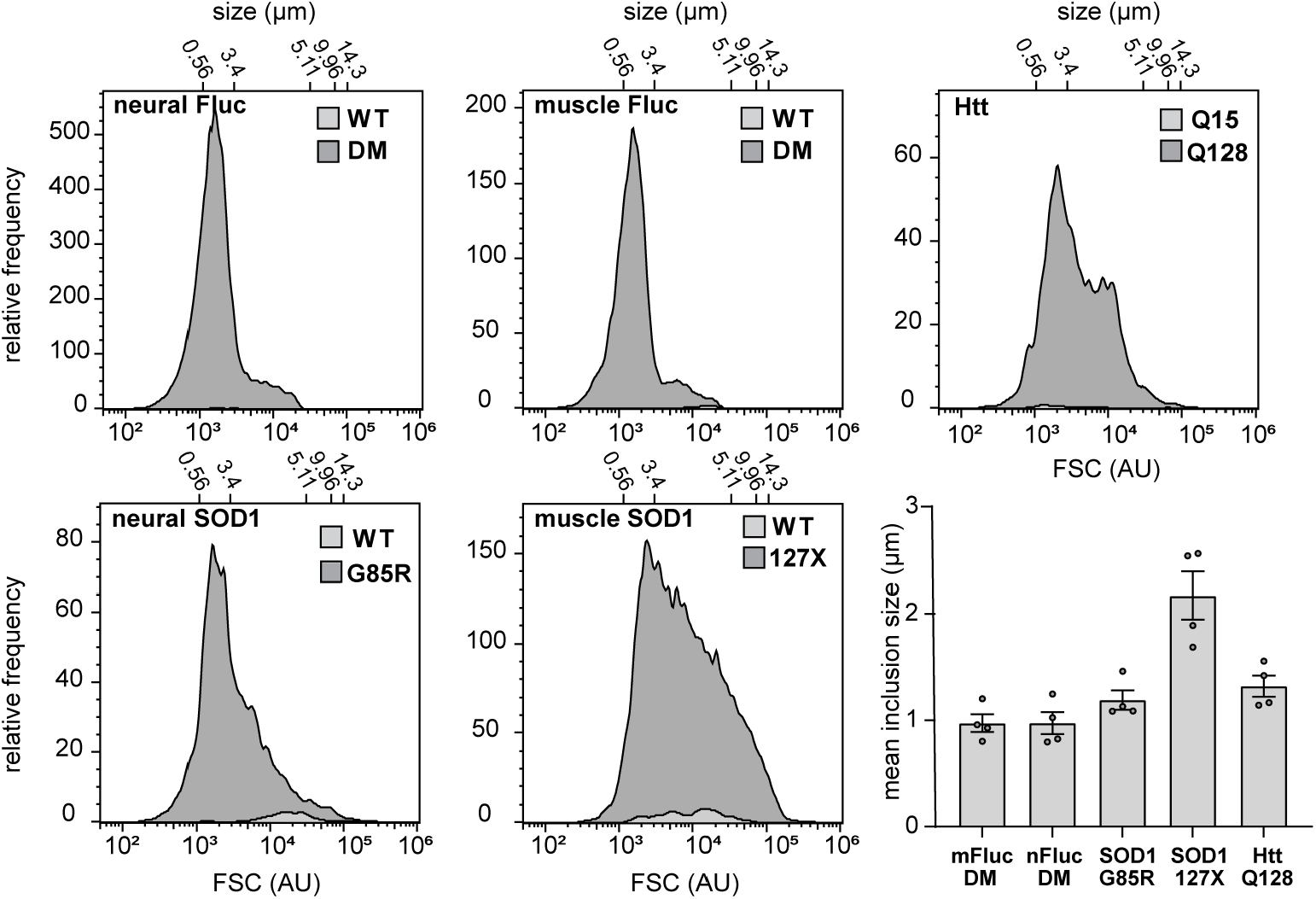
Flow cytometry can be used to determine the apparent size of protein inclusions in *C. elegans* lysate. Frequency histograms showing the distribution of FSC (corresponding sizes in μm indicated at top of each graph) of inclusions detected by flow cytometry in the *C. elegans* strains used in this study. Note that in most cases the signal from WT SOD1, FlucWT and HttQ15 is too small to be observable in these plots. Quantification of the mean inclusion size for each aggregation-prone strain is shown (*bottom right panel*). Values are means ± S.E.M from 4 independent replicates. Data from Fluc-expressing strains represent worms that were heat shocked (at 35°C for 1 h) to induce aggregation prior to immediate lysis and analysis.

## Discussion

Recently developed flow cytometry approaches for quantifying protein aggregation in cellular model systems have shown great promise in providing a simple and rapid alternative to traditional methods (Hidalgo et al. 2016; McMahon et al. 2021; Ramdzan et al. 2012; Shiber et al. 2014; Whiten et al. 2016). In this study, we describe a method that enables fluorescent inclusions to be enumerated and characterised in *C. elegans* by exploiting a standard flow cytometry setup. This method relies on mechanically lysing worms through sonication, identifying inclusions based on their fluorescence, and then normalising this number to the concentration of protein in the lysate. We demonstrate that this technique is robust and versatile: it can be applied to multiple *C. elegans* strains that model aggregation by expressing proteins of interest, including those associated with neurodegenerative diseases such as ALS and Huntington’s disease, in muscle or neural cells. We further validated this flow cytometry-based technique by demonstrating that dynamic changes in inclusion formation could be detected when known modulators of protein homeostasis were applied. Namely, the robust increase in the number of FlucDM inclusions in response to acute heat stress (and subsequent decrease during a recovery period) could be quantified in strains expressing this protein in neurons and body-wall muscle cells. Evident from both our imaging and flow cytometry data, we found that while the number of FlucDM inclusions in both strains decreased after a period of recovery, the recovery in the neural-expressing strain was more efficient than in the muscle-expressing strain. This is in accord with the findings of Gupta et al. (2011), who reported a similar observation in young FlucDM-expressing worms. This result highlights the applicability of our technique for the study of the aggregation process and how it differs across tissues in *C. elegans*. Together, these results demonstrate the applicability of our technique for the study of the protein inclusions in *C. elegans*, including modulators of their formation.

Previous approaches for analysing whole worms using large particle flow cytometers have been reported, including the Complex Object Parametric Analyser and Sorter (COPAS). This has allowed some high-throughput analysis for scoring various worm phenotypes on the basis of length, optical density and fluorescence (Pulak 2006). However, this approach is expensive and has limitations in its applicability to quantify aggregation in worms as individual inclusions are not detectable. Thus, applications of sorting by fluorescence are mainly limited to studies of gene reporter expression. By analysing the lysate from worms, our flow-cytometry based approach is based on the same principle that underpins techniques such as the filter-trap assay, i.e. as the protein of interest aggregates into inclusions, it transitions from a soluble to insoluble state. Moreover, our flow cytometry-based method is especially powerful with regards to quantifying and characterising the range of inclusions that can form *in vivo* in worms. Notably, even small inclusions (~0.5 μm) could be (rapidly) quantified, something that can be challenging with microscopy as a high level of magnification is required to resolve them (Link et al. 2006; Wang et al. 2009). For example, we can quantitatively measure inclusion formation in FlucDM-expressing worm strains using flow cytometry, whereas quantification of these inclusions using fluorescence microscopy would be difficult due to the spatial overlap of inclusions observed in images. Other biochemical techniques, such as immunoblotting of fractionated lysates or filter trap assays, can overcome some of the limitations of microscopy imaging and can provide a quantitative analysis of the amount of insoluble protein in a sample. However, these techniques are time-consuming and laborious, and often require a degree of optimisation. Another major advantage of exploiting flow cytometry to quantify these inclusion is that the lysate from worms can be analysed directly (within minutes of lysis) and can be used to determine other physicochemical properties of the inclusions that are formed, such as their apparent size.

One limitation with our method is that mechanical disruption is required to lyse the worms due to their thick outer cuticle. Although the sonication time used to lyse worms in this study was optimised to be as short as possible, to allow efficient worm lysis while preventing mechanical disruption of inclusions, we note that sonication may cause fragmentation of large inclusions. However, by exploiting the linear correlation between forward scatter and particle size of the flow cytometer we were able to quantify the size of inclusions in the lysates and found that the smaller, more compact aggregates were not significantly affected by sonication, as they aligned well with their apparent size *in vivo*, as observed via fluorescence micrographs.

In summary, our technique provides a simple, rapid and powerful unbiased alternative to traditional methods of quantifying inclusions in a broad range of *C. elegans* models of protein aggregation. Our method builds upon the previously reported FloIT technique (Whiten et al. 2016) by extending it for the analysis of inclusions formed in an organism. It can be used to complement other methods of quantifying aggregation, such as imaging and filter-trap assays. Moreover, our method has the potential to be applied to medium- and high-throughput screens to identify genes that modulate aggregation or compounds that reduce inclusion formation in *C. elegans* models of protein aggregation.

## Materials and methods

### C. elegans strains and growth conditions

*C. elegans* strains: wildtype (N2 Bristol), EAK102 [*eeeIs1(unc-45p::Htt513(Q15)::YFP::unc-45 3’UTR)*], EAK103 [*eeeIs2(unc-45p::Htt513(Q128)::YFP::unc-45 3’UTR)*], IW31 [*(iwIs27(snb-1p::hSOD1-WT::YFP)*], IW8 [*iwIs8(snb-1p::hSOD1-G85R::YFP)*], AM263 [*rmIs175(unc-54p::hSOD1-WT::YFP)*], AM725 [*rmIs290(unc-54p::hSOD1-127X::YFP)*] were obtained from the *Caenorhabditis* Genetics Centre (University of Minnesota). The strains: FUH62 [*rol-6(su1006); marIs62(unc-54p::Fluc-WT::EGFP)*], FUH135 [*rol-6(su1006); marIs135(unc-54p::Fluc-DM::EGFP)*], FUH48 [*rol-6(su1006); marIs48(F25B3*.*3p::Fluc-WT::EGFP)*], FUH137 [*rol-6(su1006); marIs137(F25B3*.*3p::Fluc-DM::EGFP)*] were a kind gift from Prof. F. Ulrich Hartl (Max Planck Institute of Biochemistry, Munich). All *C. elegans* strains used in this study were cultured on nematode growth medium (NGM) plates seeded with the OP50 *E. coli* strain, according to standard methods (Brenner 1974). Animals were kept at 20°C unless otherwise indicated. Larval stage 4 (L4) is considered day 0 for all experiments. All other materials used in this study were obtained from Thermo Fisher Scientific, unless otherwise stated.

### Worm synchronisation and sample preparation

Age synchronisation of worms was performed by preparing 3 NGM plates per strain by placing ~30 young (day 1-2) adult hermaphrodites on each plate and allowing them to lay eggs for a period of 4 h such that ~500 synchronous eggs of each strain were produced. The gravid adults were then removed from the plates leaving the age synchronised eggs. The plates were then incubated at 20°C until the hatched larvae were at the day 1 adult stage (~72 h). Adult nematodes were then washed off each plate with M9 buffer (22 mM KH_2_PO_4_, 42 mM Na_2_HPO_4_, 85.5 mM NaCl, 1 mM MgSO_4_, pH 7.0). Worms were placed in 1.5 mL microfuge tubes and left for 2 min to allow the worms to settle into a pellet at the bottom of the tubes. After removing the supernatant, worm pellets were washed once with fresh M9 buffer to remove residual OP50 bacteria, and the supernatant was once again removed from the worm pellets. Worm pellets were then resuspended in NP-40 lysis buffer (50 mM Tris-HCl, 150 mM NaCl, 1 mM EDTA, 1% (v/v) Nonidet™ P-40, supplemented with 0.5% (v/v) Halt™ Protease Inhibitor Cocktail, pH 8.0). The resuspended samples were then lysed by pulse sonication using a Sonifer^®^ 250 Digital cell disruptor with a double step micro-tip (Branson Ultrasonics, Brookfield, CT, USA) set to 40% amplitude (1 s pulses followed by 30 s rest for a total sonication time of 5 s). Worm lysates were then clarified by centrifuging (2 min; 4°C; 500 × *g*) to remove unlysed worms and other large debris. The clarified supernatant was then placed in flow cytometry tubes and kept on ice prior to analysis by flow cytometry.

For experiments assaying worms at different ages, NGM plates containing 50 μM 5-fluorodeoxyuridine (FUDR) were used instead of standard NGM plates to inhibit egg-laying, as described previously (Sutphin & Kaeberlein 2009). In these experiments, worms were transferred to fresh FUDR-containing plates whenever OP50 bacteria were close to being depleted (approximately every 3 days). Worms were harvested from the plates when they reached the required age using the same procedure as above, snap frozen in liquid nitrogen, and stored at −20°C until ready for use. For experiments involving heat shock, plates of worms were placed in a conventional air incubator (Thermoline, TI-20F) at the temperature indicated in the figure.

### Imaging of transgenic worms

Synchronised day 1 adult worms were anaesthetised using 100 mM sodium azide and mounted on 2% agarose pads and covered with a glass coverslip for imaging. Images were taken using either a Leica SP8 FALCON laser scanning confocal microscope or a Leica DMi8 fluorescence microscope (Leica Microsystems, Wetzlar, Germany). EGFP and YFP molecules were illuminated by excitation at 488 nm and the SP8 FALCON was set to detect between 498 and 550 nm using a photomultiplier tube detector with laser power was set to 20%. All images were taken using either a 20X dry objective or a 63X oil immersion objective. In each image, at least 15 Z-stacks were taken and the final image used was a Z-stack projection. Following acquisition, images were prepared using the Leica Application Suite X (LAS X) software (Leica Microsystems).

### Analysis of worm lysates by flow cytometry

Worm lysate was analysed using an Attune NxT flow cytometer. Forward scatter (FSC), side scatter (SSC) and GFP/YFP fluorescence (488 nm excitation, 530/30 collection) were the parameters measured with the axes set to log10. For all experiments, the voltages were set to 40 (FSC), 340 (SSC) and 320 (GFP/YFP) arbitrary units and the threshold for detected particles was set to 100 (FSC) and 100 (GFP/YFP) arbitrary units, which was the lowest possible threshold for these parameters in order to detect small inclusions. The flow cytometer was set to analyse 50 μL of each sample using a flow rate of 100 μL/min. After flow cytometric analysis, the total protein concentration of each sample was determined by performing a BCA assay using the remaining lysate, according to the manufacturer’s instruction. The FCS files generated by flow cytometry were analysed using FlowJo version 10 (Tree Star Ashland, OR, USA). Inclusions were detected based on their FSC and GFP/YFP signal. Non-fluorescent N2 control worm lysate was used in each experiment to assist with identifying and gating inclusions. The number of inclusions per μg of protein in the lysate (*i*) was determined using the following formula:

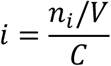

Where *n*_*i*_ is the number of inclusions detected, *V* is the volume of lysate analysed, and *C* is the concentration of the sample in μg/μL.

### Immunoblotting

For immunoblotting experiments, worms were synchronised, harvested and lysed as described above (*Worm synchronisation and sample preparation*), with the exception that 6 plates of synchronised worms were used for each strain instead of 3, to increase total protein yield. Once lysates were obtained, the total protein concentration for each sample was determined using a BCA assay, and each sample was adjusted with NP-40 lysis buffer such that they all had equal concentrations of total protein (1 mg/mL). An aliquot (30 μL) of each sample was used as a loading control for SDS-PAGE electrophoresis and kept on ice. The remaining sample was centrifuged at 20 000 × *g* for 20 min at 4°C. The supernatant (NP-40 soluble fraction) was then carefully collected and kept on ice. The remaining protein pellet was washed once with ice-cold TNE buffer (50 mM Tris-HCl, 150 mM NaCl, 1 mM EDTA, pH 8.0), and centrifuged again (20 min; 4°C; 20 000 × *g*). The supernatant was then carefully removed, and the protein pellet (insoluble fraction) was resuspended in 50 μL urea buffer (7M Urea, 50 mM Tris-HCl, 4% CHAPS (w/v), pH 7.5) to solubilise the protein for immunoblotting. SDS-PAGE loading buffer (final concentrations: 500 mM Tris-HCl, 2% (w/v) SDS, 25% (w/v) glycerol, 0.01% (w/v) bromophenol blue, 15% (v/v) β-mercaptoethanol (Sigma-Aldrich), pH 6.8) was added to each sample followed by heating at 95°C for 5 min.

Samples were loaded onto Any kD Mini-PROTEAN TGX Stain-Free Protein Gels (Bio-Rad, Hercules, CA, USA) with Precision Plus Protein™ dual colour molecular weight standards (Bio-Rad) used as protein size markers. Acrylamide gels were electrophoresed in SDS-PAGE running buffer (192 mM glycine, 3.5 mM SDS, 25 mM Tris(hydroxymethyl)methylamine) at 150 V for ~ 60 min, or until the bromophenol blue dye front reached the bottom of the resolving gels. Following electrophoresis, stain-free gels were imaged using a Gel Doc™ EZ Imager (Bio-Rad) (in order to confirm that the relative amount of total protein loaded into the wells was similar among samples) and then proteins on each gel were transferred to an ImmunoBlot™ polyvinylidene difluoride (PVDF; Bio-Rad) membrane by running at 100 V for 1 h in ice-cold transfer buffer (25 mM Tris base, 192 mM glycine, 20% (v/v) methanol, pH 8.3). After the transfer, PVDF membranes were blocked with 5% (w/v) skim milk powder in Tris-buffered saline (TBS; 50 mM Tris base, 150 mM NaCl, pH 7.6) at 4°C overnight. Membranes were then incubated with rabbit anti-GFP (1:2500; ab290, Abcam, Cambridge, MA, USA) diluted in 5% (w/v) skim milk powder in TBS-T (TBS with 0.05% Tween-20) for 2 h at room temperature (22°C). Membranes were washed four times (each for 10 min) in TBS-T before being incubated with an anti-rabbit horse radish peroxidase (HRP)-conjugated secondary antibody (31466, Thermo Fisher Scientific), diluted 1:5000 into 5% (w/v) skim milk powder in TBS-T. Membranes were rocked at room temperature for 1 h before being washed four times (each for 10 min) in TBS-T. The EGFP/YFP tags on the proteins was detected by applying SuperSignal^®^ West Dura Extended Duration Chemiluminescent Substrate and imaging with an Amersham Imager 600RGB (GE Healthcare Life Sciences, Little Chalfont, UK).

### Statistical analysis

Graphs were generated from data acquired from flow cytometry experiments using GraphPad Prism version 9 (GraphPad Software, San Diego, CA, USA). Data are reported as mean ± standard error of the mean (S.E.M). Statistical analyses on the flow cytometry data were also performed using the GraphPad software by either one-way or two-way analysis of variance (ANOVA) and, where appropriate, a Tukey’s post-hoc test. Where two sets of variables were compared, data were analysed by a two-tailed student t-test, assuming equal variance. An alpha level of 0.05 was considered statistically significant in all analyses.

## Acknowledgments

We are grateful to the technical staff at Molecular Horizons and the Illawarra Health and Medical Research Institute for technical and administrative support. We thank Prof. F. Ulrich Hartl (Max Planck Institute of Biochemistry, Munich) for providing the FUH62, FUH135, FUH48 and FUH137 strains, and the *Caenorhabditis* Genetics Centre, which is supported by the National Institutes of Health (P40 OD010440), for providing the rest of the strains used in this study.

